# Metabolic versatility of the nitrite-oxidizing bacterium *Nitrospira marina* and its proteomic response to oxygen-limited conditions

**DOI:** 10.1101/2020.07.02.185504

**Authors:** Barbara Bayer, Mak A. Saito, Matthew R. McIlvin, Sebastian Lücker, Dawn M. Moran, Thomas S. Lankiewicz, Christopher L. Dupont, Alyson E. Santoro

## Abstract

The genus *Nitrospira* is the most widespread group of chemolithoautotrophic nitrite-oxidizing bacteria that thrive in diverse natural and engineered ecosystems. *Nitrospira marina* Nb-295^T^ represents the type genus and was isolated from the oceanic water column over 30 years ago, however, its genome has not yet been analyzed. Here, we analyzed the complete genome sequence of *N. marina* and performed select physiological experiments to test genome-derived hypotheses. Our data confirm that *N. marina* benefits from additions of undefined organic carbon substrates, has adaptations to combat oxidative, osmotic and UV-light induced stress and low dissolved *p*CO_2_, and is able to grow chemoorganotrophically on formate. We further investigated the metabolic response of *N. marina* to low (∼5.6 µM) O_2_ concentrations commonly encountered in marine environments with high nitrite concentrations. In response to O_2_-limited conditions, the abundance of a potentially more efficient CO_2_-fixing pyruvate:ferredoxin oxidoreductase (POR) complex and a high affinity *cbb3*-type terminal oxidase increased, suggesting a role in sustaining nitrite oxidation-driven autotrophy under O_2_ limitation. Additionally, a Cu/Zn-binding superoxide dismutase increased in abundance potentially protecting this putatively more O_2_-sensitive POR complex from oxidative damage. An increase in abundance of proteins involved in alternative energy metabolisms, including type 3b [NiFe] hydrogenase and formate dehydrogenase, indicate a high metabolic versatility to survive conditions unfavorable for aerobic nitrite oxidation. In summary, the genome and proteome of the first marine *Nitrospira* isolate identifies adaptations to life in the oxic ocean and provides important insights into the metabolic diversity and niche differentiation of NOB in marine environments.

## Introduction

Aerobic nitrite (NO_2_^-^) oxidation is the main biochemical nitrate (NO_3_^-^)-forming reaction, carried out during the second step of nitrification [1]. In marine ecosystems, nitrate is the dominant form of biologically available nitrogen, which is rapidly assimilated by phytoplankton in surface waters and accumulates in the deep sea [2].

Nitrite-oxidizing bacteria (NOB) are chemolithoautotrophic microorganisms comprising seven known genera (*Nitrobacter, Nitrococcus, Nitrospina, Nitrospira, Nitrotoga, Nitrolancea*, and *Candidatus* Nitromaritima) in four phyla [3]. The genus *Nitrospira* consists of at least six phylogenetic sublineages and is the most diverse NOB genus [3]. *Nitrospira* are ubiquitously present in natural and engineered ecosystems, including oceans [4, 5], freshwater habitats [6], soils [7, 8], saline-alkaline lakes [9], hot springs [10], wastewater treatment plants [11–13], and aquaculture biofilters [14, 15]. In human-made ecosystems, *Nitrospira* is generally considered to be adapted to low NO_2_^-^ concentrations [16]. In the open ocean, however, where NO_2_^-^ concentrations are exceedingly low, NOB affiliated with the phylum *Nitrospinae* appear to be the dominant nitrite oxidizers [4], whereas *Nitrospira* bacteria are found in relatively high NO_2_^-^ environments such as sediments and deep sea hydrothermal vent plumes [17, 18]. *Nitrospira* also dominate over *Nitrospinae*-affiliated bacteria in some deep-sea trench environments [19, 20]. High NO_2_^-^ concentrations are found coincident with low O_2_ concentrations in oxygen minimum zone (OMZ) waters [21, 22] in a feature known as the secondary nitrite maximum. Despite the O_2_-dependency of all known NOB, NO_2_^-^ oxidation can still be detected at nanomolar O_2_ concentrations [23].

Some metabolic features appear to be common among *Nitrospira*, based on genomic analyses to date. Metagenomic analysis of *Ca*. Nitrospira defluvii, a representative of *Nitrospira* sublineage I, indicated a periplasmic location of nitrite oxidoreductase (NXR), the key enzyme of the NO_2_^-^ oxidation, and the presence of the O_2_-sensitive reductive tricarboxylic acid (rTCA) cycle for carbon fixation [24]. These results suggest that *Nitrospira* evolved from microaerophilic or anaerobic ancestors [24]. *Ca*. N. defluvii lacks genes for classical oxidative stress defense enzymes present in most aerobic organisms including catalase and superoxide dismutase, indicative for adaptations to low O_2_ environmental niches [24]. However, other *Nitrospira* species encode both types of enzymes [25, 26], suggesting different O_2_ tolerances within members of the *Nitrospira* genus. In addition to NO_2_^-^ oxidation, *Nitrospira* bacteria have been shown to exhibit a high metabolic versatility, growing aerobically on hydrogen [27] or anaerobically on organic acids while respiring nitrate [25]. Recently, the capability for the complete oxidation of ammonia to nitrate (comammox) was identified in representatives of sublineage II *Nitrospira* [14, 28], however, comammox *Nitrospira* appear to be absent in marine systems [28].

*N. marina* Nb-295^T^, the type species of the genus *Nitrospira*, was isolated from a water sample collected at a depth of 206 m from the Gulf of Maine in the Atlantic Ocean over 30 years ago [5]. It is the only *Nitrospira* species isolated from the oceanic water column, however, its genome has not yet been analyzed. Here, we analyzed the complete genome sequence of *N. marina* Nb-295^T^ and compared the proteome signatures of cultures grown under atmospheric O_2_ tension and under low O_2_ conditions.

## Materials and Methods

### Cultivation of *N. marina* Nb-295

*Nitrospira marina* Nb-295^T^ was obtained from the culture collection of John B. Waterbury at the Woods Hole Oceanographic Institution (WHOI). To test the effect of different organic substrates on strain Nb-295, cultures were grown in 50 ml polycarbonate bottes in autotrophic mineral salts medium at pH 7.8 containing 2 mM NaNO_2_ (Table S1), and bottles were incubated at 25°C in the dark without agitation. The following organic carbon substrates were individually added to the culture medium of duplicate cultures: 150 mg L^-1^ yeast extract, 150 mg L^-1^ tryptone, 1 mM pyruvate, 1 mM formate, or 1 g L^-1^ glycerol. NO_2_^-^ consumption was measured as previously described [29] and growth was monitored by flow cytometry (see Supplementary Methods). To test for chemoorganotrophic growth, NO_2_^-^ was omitted from the culture medium.

For incubations at different O_2_ concentrations, triplicate cultures of *N. marina* Nb-295 were grown at 22°C in 400 mL of mineral salts medium containing 70% seawater and 2 mM NaNO_2_ as described by Watson *et al*. [5] in 500 mL polycarbonate bottles with a custom-made sparging rig. Bottles were constantly bubbled with one of two sterile custom gas mixes containing either 0.5% or 20% oxygen, 300 ppm CO_2_, and a balance of high-purity N_2._ NO_2_^-^ concentrations were measured as a proxy for growth as described above and 10 mL aliquot of each culture was fixed at the last time point (2% formaldehyde, 1 h, 4°C) for cell enumeration on an epifluorescence microscope as previously described [30].

### DNA extraction, genome sequencing and annotation

High molecular weight genomic DNA was extracted from stationary phase cultures using a CTAB extraction protocol [31] and sequenced on the PacBio platform at the US Department of Energy Joint Genome Institute (JGI). 754,554 reads were produced, with 209,987 passing quality control. The assembly was conducted using HGAP (v 2.2.0.p1) with improvement with Quiver [32] resulting in a single contig.

Gene annotation was conducted using JGI’s Integrated Microbial Genomes and Microbiomes (IMG/M) pipeline [33] and the MicroScope platform [34]. Manual curation included sequence similarity searches using BLASTP [35] against the Transporter Classification database [36] and protein domain searches using InterProScan (release 72.0) [37]. Signal peptides were identified with SignalP 5.0 [38] to determine if proteins were potentially addressed to the membrane and/or released to the periplasmic space. A list of manually curated annotations can be found in Table S5.

Phylogenomic analysis was performed using 120 concatenated phylogenetic marker genes of representatives of the phylum *Nitrospirae/Nitrospirota* as implemented in the Genome Taxonomy Database Toolkit (GTDB-tk) version 1.1.1. [39] (see Supplementary Methods).

### Protein extraction and proteome analyses

Cells were harvested for proteomic analysis during exponential growth when [NO_2_^-^] dropped to ∼ 500 µM. Each culture was mixed with an equal volume of a house-made fixative [40] similar to the commercially available solution RNALater (Thermo Fisher), and filtered by vacuum filtration onto 25 mm, 0.2 µm pore size Supor filters (Pall). 200 mL of fixed culture (equivalent to 100 mL of growth medium) were filtered for protein extraction and proteomic analysis. Filters were frozen at −80°C until extraction. A detailed description of the procedures used for protein extraction and purification can be found in the Supplementary Methods. Procedures for mass spectrometry analyses can be found in Saito *et al*. [41]. In addition, targeted quantitative proteomics using custom-made isotopically-labelled (^15^N) peptide standards was performed as previously described [41]. Tryptic peptides from different proteins, including nitrite oxidoreductase (NXR), were targeted for absolute quantitation (see Table S2). Cellular NXR concentrations were calculated based on NxrA peptide concentrations using a cellular carbon content of 152 fg cell^-1^ (Santoro et al, *unpublished*) and an estimated cellular protein content of 50% (Table 1). NXR complex density on the cellular membrane was calculated using an estimated NXR complex size of 63 nm^2^ [42] and an estimated surface area of 2.14 µm^2^ (assuming a cylindrical shape with a length of 1.5 µm and a radius of 0.2 µm).

**Table 1.** Quantitative proteomics of selected proteins using isotopically-labelled (^15^N) peptide standards. Nitrite oxidoreductase peptide targets were designed to match a specific NxrA copy (1, 2 or 3), or to match all three as indicated in the target protein column. Values represent the mean of triplicate measurements and standard deviations are shown in brackets.

| Target protein* | $^{15}\text{N}$-Peptide sequence | Peptide concentration (copies cell$^{-1}$) | |
| --- | --- | --- | --- |
|  |  | atm. O<sub>2</sub> | low O<sub>2</sub> |
| Nitrite oxidoreductase, alpha subunit 1 | LVVITPEYNPTAYR | **845 (52)** | **1,924 (275)** |
|  | DYAFPDFANSYSGK | **241 (40)** | **451 (91)** |
|  | HPFWEETNESKPQWTR | **673 (39)** | **1,571 (388)** |
| Nitrite oxidoreductase, alpha subunit 2 | VVVITPEYNPTAQR | 1,154 (27) | 851 (187) |
| Nitrite oxidoreductase, alpha subunit 3 | DYQFPDFTSTYSGK | 1,761 (159) | 1,261 (242) |
|  | SGIDPALTGTHR | 4,519 (516) | 3,400 (526) |
|  | IAVITPEYNPTAYR | 4,860 (496) | 3,741 (710) |
| Nitrite oxidoreductase, alpha subunit (all) | GWKPSDPYYK | 11,467 (1,957) | 9,564 (1,083) |
|  | AIALDTGYQSNFR | 13,996 (962) | 13,116 (2,934) |
| Pyruvate:ferredoxin oxidoreductase,<br>alpha subunit (PorA) | TPSFFTGSEVIK | 2,026 (192) | 1,550 (248) |
|  | EAIAILEEEGIR | 1,362 (80) | 1,054 (170) |
|  | EVSATVPNNER | 2,453 (169) | 1,736 (215) |
| Ribonucleotide reductase | TGESPYQTIPFSHR | 300 (46) | 198 (30) |
|  | EAAVPEPYIHR | 705 (132) | 472 (77) |
|  | IINQSLPPALR | 750 (88) | 546 (68) |

\*A complete list of quantified proteins can be found in Supplementary table 2.

Differential levels of expression between the two treatments (i.e., atmospheric O_2_ and low O_2_ conditions) were tested with the DESeq2 Bioconductor package (version 1.20.0) [43] in the R software environment (version 3.5.0) [44] using spectral counts as input data as previously described [45]. Proteins with a mean spectral count below 6 across all treatments were excluded from the analysis. In DESeq2, only proteins that increased in abundance under low O_2_ conditions were considered (‘altHyphothesis=greater’), P values were adjusted using the Benjamini-Hochberg method (‘pAdjustmethod=BH’) and independent filtering was omitted (‘independentFiltering=FALSE’). Changes in protein abundance were considered statistically significant when adjusted P values were lower than or equal to 0.05 (see Table S3). While DESeq2 has a high precision and accuracy [46], it is more conservative than other methods on low-count transcripts/proteins [47]. Protein abundances were visualized with the pheatmap package (version 1.0.12) [48] in the R software environment [44]. The normalized spectral abundance factor (NSAF) was calculated as proxy for relative protein abundances [49], and values were square-root transformed to improve visualization of low abundant proteins.

## Results and Discussion

### Genome analysis

The genome of *N. marina* Nb-295 is a single element of 4,683,627 bp and contains 4272 coding DNA sequences (CDS) including one rRNA operon. A 5578 bp region of 99.7% identity on each end of the scaffold suggests circularization into a single chromosome. No plasmids or extra-chromosomal elements were identified. The G+C content is 50.04%, which is lower than other *Nitrospira* species and the marine nitrite-oxidizers *Nitrospina gracilis* and *Nitrococcus mobilis* (Table S4). Phylogenomic analysis of available closed genomes, metagenome-assembled genomes (MAGs) and single-amplified genomes (SAGs) affiliated with the phylum *Nitrospirae/Nitrospirota* (see Supplementary Methods) placed *N. marina* Nb-295 within a cluster of genomes derived from marine and saline environments (Fig. 1). The most closely related *Nitrospira* MAG (UBA8639) was obtained from a laboratory-scale nitrification reactor, however, the reactor influent consisted to 33% of untreated seawater [50], suggesting a marine origin of this MAG. The 16S rRNA gene sequence of *N. marina* Nb-295 clustered together with environmental sequences derived from marine sediments and marine aquaculture biofilters (Fig. S1), and shared 99.1% and 97.9% sequence identity with the cultured lineage IV representatives *Nitrospira sp*. Ecomares 2.1 [15] and *Ca*. Nitrospira alkalitolerans [9], respectively.

**Fig 1.**
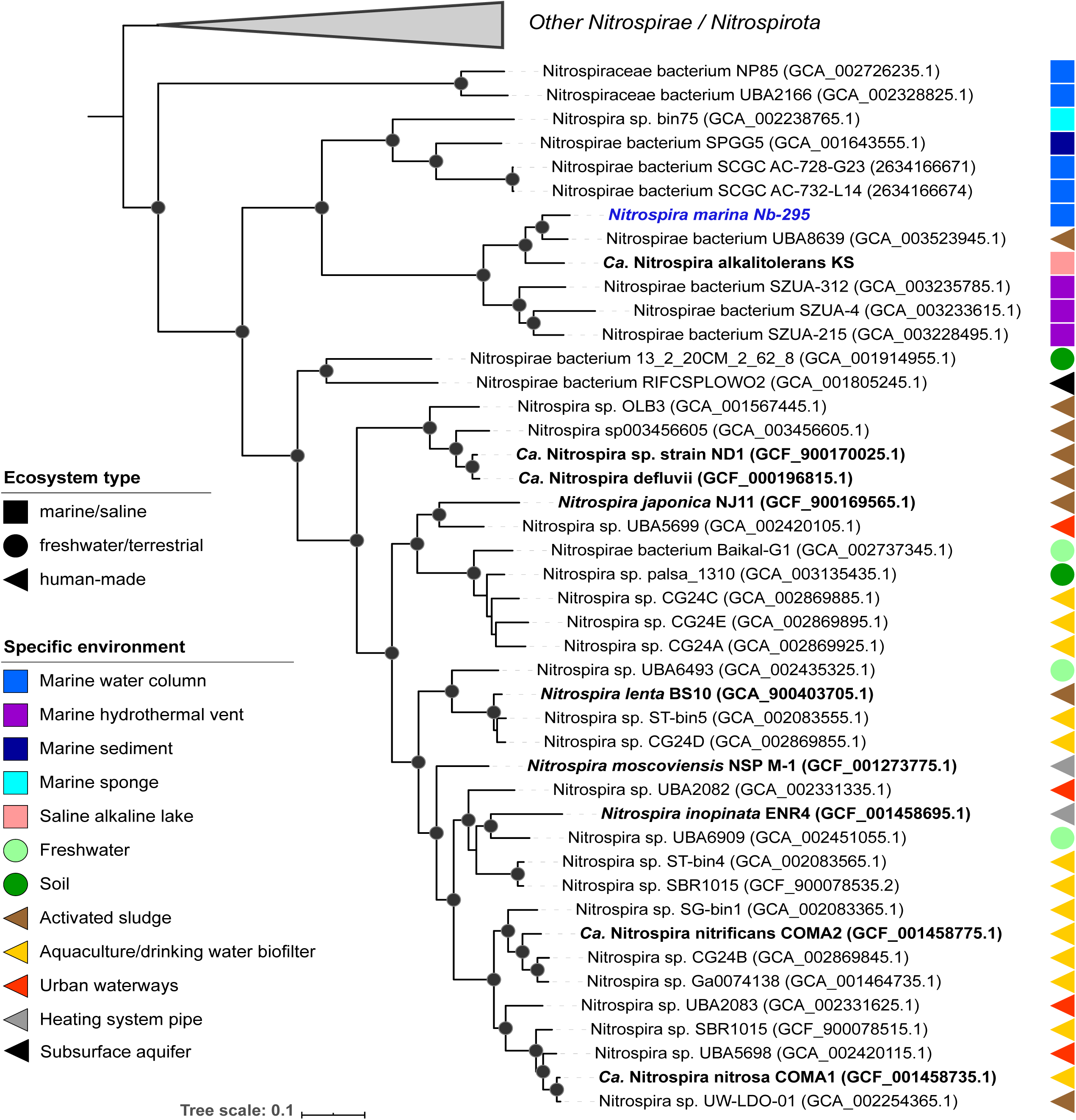
Maximum likelihood phylogenetic tree of representatives of the phylum *Nitrospirae/Nitrospirota*. The multiple sequence alignment consisting of 120 concatenated phylogenetic marker genes contained 95 genomes and metagenome-assemble genomes (MAGs) from the Genome Taxonomy Database (GTDB) (Release 04-RS89, 19 June 2019), the genome of *N. marina*, the MAG of *Ca*. N. alkalitolerans and two open ocean SAGs AC-738-G23 and AC-732-L14. All MAGs and SAGs were estimated to be ≥50% complete with ≤5% contamination. Nodes with UFBoot support of at least 95% are indicated as black filled circles. The scale bar represents 0.1 substitutions per site.

#### Nitrogen metabolism and respiratory chain

The genome of *N. marina* encodes orthologs of all known proteins required for NO_2_^-^ oxidation, including the putatively membrane-associated periplasmic nitrite oxidoreductase (NXR). Three candidates each of NxrA and NxrB, three putative NxrC candidates and two NxrC-like proteins were identified in the *N. marina* genome (Table S4 and S5). The genes *nxrA* and *nxrB*, encoding the alpha and beta subunits of NXR, are co-localized in three clusters whereas all *nxrC* candidate genes are localized separately from *nxrAB*, as previously described for *N. moscoviensis* [51]. The NxrA subunits share 87.3-88.9% amino acid identity, whereas the NxrB subunits share 98.8-99.5% amino acid identity with each other. The putative NxrC subunits are less conserved, sharing between 33.7.7-86.9% amino acid identity.

Like all other analyzed *Nitrospira* genomes, *N. marina* encodes a putative copper-dependent NO-forming nitrite reductase (NirK), yet its function in *Nitrospira* and other NOB remains unknown [52]. *N. marina* also encodes the ferredoxin-dependent nitrite reductase (NirA) for assimilatory nitrite reduction, which appears to be conserved in *Ca*. Nitrospira lenta and *Ca*. N. defluvii but absent in other *Nitrospira* species [52]. Additionally, *N. marina* encodes three high affinity ammonium transporters (Amt) enabling direct uptake of reduced N for assimilation, and cyanate lyase to hydrolyze cyanate to ammonium and CO_2_ (Table S5). In contrast to some *Nitrospira* species [13, 25, 28], *N. marina* does not encode a urease operon or a urea transporter, which would catabolize urea to ammonia for N assimilation (Table S4).

Previous genome studies of *Nitrospira* revealed multiple copies of several complexes of the respiratory chain [24–26, 51]. *N. marina* encodes two paralogous copies of complex I, one of which contains a duplication of NuoM and lacks genes for NuoE, NuoF and NuoH (Table S5), which is a characteristic feature of *Nitrospira* genomes [51, 53]. Further, the *N. marina* genome contains two copies of complex III, two cytochrome *bd* oxidases and seven putative cytochrome *bd*-like oxidases (Table S5), which show limited partial similarity to canonical cytochrome *bd* oxidase as described for *N. moscoviensis* [51]. One of these cytochrome *bd*-like oxidases (*bd*-like_6) contains putative heme b and copper binding sites potentially functioning as a novel terminal oxidase as previously proposed [24]. Additionally, *N. marina* encodes for a *cbb*_3_-type terminal oxidase, which usually exhibit high affinities for O_2_ [54]. This feature is shared with the closely related *Ca*. N. alkalitolerans [9] and with the more distantly related marine NOB *Nitrospina gracilis* [55], but absent in all other thus far sequenced *Nitrospira* species. In addition to the canonical H^+^-translocating F_1_F_O_-ATPase (complex V), *N. marina* also encodes a putative alternative Na^+^-translocating N-ATPase (Table S5), which potentially contributes to the maintenance of the membrane potential and the generation of a sodium motive force (SMF) as suggested for *Ca*. N. alkalitolerans [9]. Furthermore, a H^+^-translocating pyrophosphatase (H^+^-PPase) with homology to *Leptospira*/protozoan/plant-type enzymes [56] was identified. H^+^-PPases couple the translocation of H^+^ to the hydrolysis of the biosynthetic by-product pyrophosphate (PP_i_), which is suggested to be an adaptation to life under energy limitation [57]. The *N. marina* genome also contains an alternative complex III (ACIII) module, which shares similarity with that from sulfur-reducing Acidobacteria [58]. Like canonical complex III, ACIII also functions as a quinol oxidase transferring electrons to cytochrome *c* and contributes to energy conservation (Refojo *et al*., 2012). With the exception of the comammox bacterium *Ca*. Nitrospira nitrificans, no homologues of ACIII modules were identified in any other NOB genome.

*N. marina* encodes a putative *hyb*-like operon containing four subunits of a cytoplasmic type 3b [NiFe] hydrogenase and six accessory proteins involved in hydrogenase assembly and maturation (Table S5). This type of hydrogenase appears to be conserved in the marine NOBs *Nitrospina gracilis* 3/211 and *Nitrococcus mobilis* Nb-231 [55, 59], *Ca*. N. alkalitolerans [9] and in comammox *Nitrospira* [14, 28] (Table S4). In addition to catalyzing the reversible, NAD^+^-dependent oxidation of hydrogen, these so-called sulfhydrogenases are able to reduce elemental sulfur (S°) or polysulfides to hydrogen sulfide (H_2_S) [60]. Furthermore, the *N. marina* genome encodes a putative periplasmic sulfite:cytochrome *c* oxidoreductase which might couple sulfite (SO_3_^2-^) oxidation to sulfate (SO_4_^2-^) with the reduction of two cytochromes as previously suggested for *Nitrospina gracilis* [55] whereas sulfide/quinone oxidoreductase, which is speculated in to mediate sulfide oxidation in *Nitrococcus* [59], is lacking. However, whether or not these enzymes are involved in energy conservation using H_2_S and SO_3_^2-^ as alternative substrates remains to be experimentally validated in NOB.

#### Central carbon metabolism

In agreement with previously characterized *Nitrospira* genomes [13, 24, 51, 52], *N. marina* encodes the complete gene repertoire for the reductive tricarboxylic acid (rTCA) cycle for carbon dioxide (CO_2_) fixation, including the key enzymes ATP-citrate lyase and 2-oxoglutarate/pyruvate:ferredoxin oxidoreductase (OGOR/POR) (Table S5). In the ocean, inorganic carbon is predominately available in the form of bicarbonate (HCO_3_^-^) and to a much lesser extent as CO_2_. Five inorganic anion transporters (SulP family) with homology to BicA HCO_3_^-^ uptake systems of the cyanobacterium *Synechococcus* [61] were identified in the *N. marina* genome (Table S5). Two of these putative BicA-like transporters are co-localized with genes encoding Na^+^/H^+^ antiporters (NhaB family), which could drive the uptake of HCO_3_^-^ via Na^+^ extrusion under alkaline conditions as suggested for the cyanobacterium *Aphanothece halophytica* [62] and *Ca*. N. alkalitolerans [9]. *N. marina* also encodes one putative SulP-related bicarbonate transporter fused to a carbonic anhydrase and four genes encoding putative alpha, beta and gamma carbonic anhydrases (Table S5), which can convert the imported HCO_3_^-^ to CO_2_ for inorganic carbon fixation via the rTCA cycle.

In addition to the rTCA cycle, *N. marina* encodes all required genes for the oxidative TCA cycle for pyruvate oxidation via acetyl-CoA, complete gluconeogenesis and glycolysis pathways, and the oxidative and non-oxidative branches of the pentose phosphate pathway (Table S5), which are common features of all sequenced *Nitrospira* genomes [9, 24–26, 52]. Furthermore, biosynthetic pathways for all amino acids except methionine were identified in the *N. marina* genome. Although *N. marina* encodes a vitamin B_12_-dependent methionine synthase (MetH) (Table S5), it appears to lack additional enzymes of known methionine biosynthesis pathways, a trait shared by all other sequenced *Nitrospira* species and *Nitrospina gracilis* [9, 24–26, 52, 55]. However, as *N. marina* can grow in artificial seawater medium without added methionine, we hypothesize that an alternative unknown pathway for the early steps of methionine biosynthesis functions in *Nitrospira and Nitrospina*. The *N. marina* genome contains genes for the biosynthesis and degradation of the storage compounds glycogen and polyphosphate (Table S5). In contrast to other *Nitrospira* and the marine NOBs *N. gracilis* and *N. mobilis* that encode a glgA-type glycogen synthase, *N. marina* encodes alpha-maltose-1-phosphate synthase (glgM) and alpha-1,4-glucan:maltose-1-phosphate maltosyltransferase (glgE) for the synthesis of glycogen via alpha-maltose-1-phosphate.

#### Use of organic substrates

*N. marina* has been reported to be an obligate chemolithotroph that grows best in medium supplemented with low concentrations of organic compounds including pyruvate, glycerol, yeast extract and peptone [5]. Thus, we investigated the genomic basis for this observation and conducted additional physiological experiments.

In addition to its complete glycolysis pathway and oxidative TCA cycle, a putative carbohydrate degradation operon was identified in the genome of *N. marina*, consisting of a sugar ABC transporter module, beta-glucosidase, a putatively secreted glycoside hydrolase (GH15) and a carbohydrate-binding protein (Table S5). *N. marina* also encodes two putative carbohydrate-selective porins (OprB), a sugar:sodium symporter (SSS family), a putative galactonate/glucarate transporter (MFS superfamily) and a putative carboxylate transporter (DASS family). Additionally, the genomic repertoire for the catabolic degradation and assimilation of peptides and amino acids, including transporter proteins for di- and oligopeptides (ABC and POT/PTR families), multiple amino acid:cation symporter (SSS, DAACS and AGCS families), amino acid/polyamine transporters (APC superfamily) and multiple putatively secreted peptidases are present in the *N. marina* genome (Table S5).

In agreement with Watson *et al*. [5], NO_2_^-^ oxidation activity was enhanced when undefined organic compound mixtures such as tryptone and yeast extract were added to the culture medium (Fig. 2). Interestingly, growth was greatly stimulated by tryptone (Fig. 2), while amendment with defined organic carbon compounds had no effect on NO_2_^-^ oxidation or growth (Table S6). Parallel incubations with ammonium as an N source did not increase activity or growth (Table S6), suggesting that the stimulating effect of tryptone, and to a lesser extent of yeast extract, most likely can be attributed to direct amino acid assimilation and does not reflect only the abolished energy demand for assimilatory NO_2_^-^ reduction. No growth on yeast and tryptone was observed in the absence of NO_2_^-^ (Fig. 3), corroborating their use as source of amino acids rather than for energy conservation.

**Fig 2.**
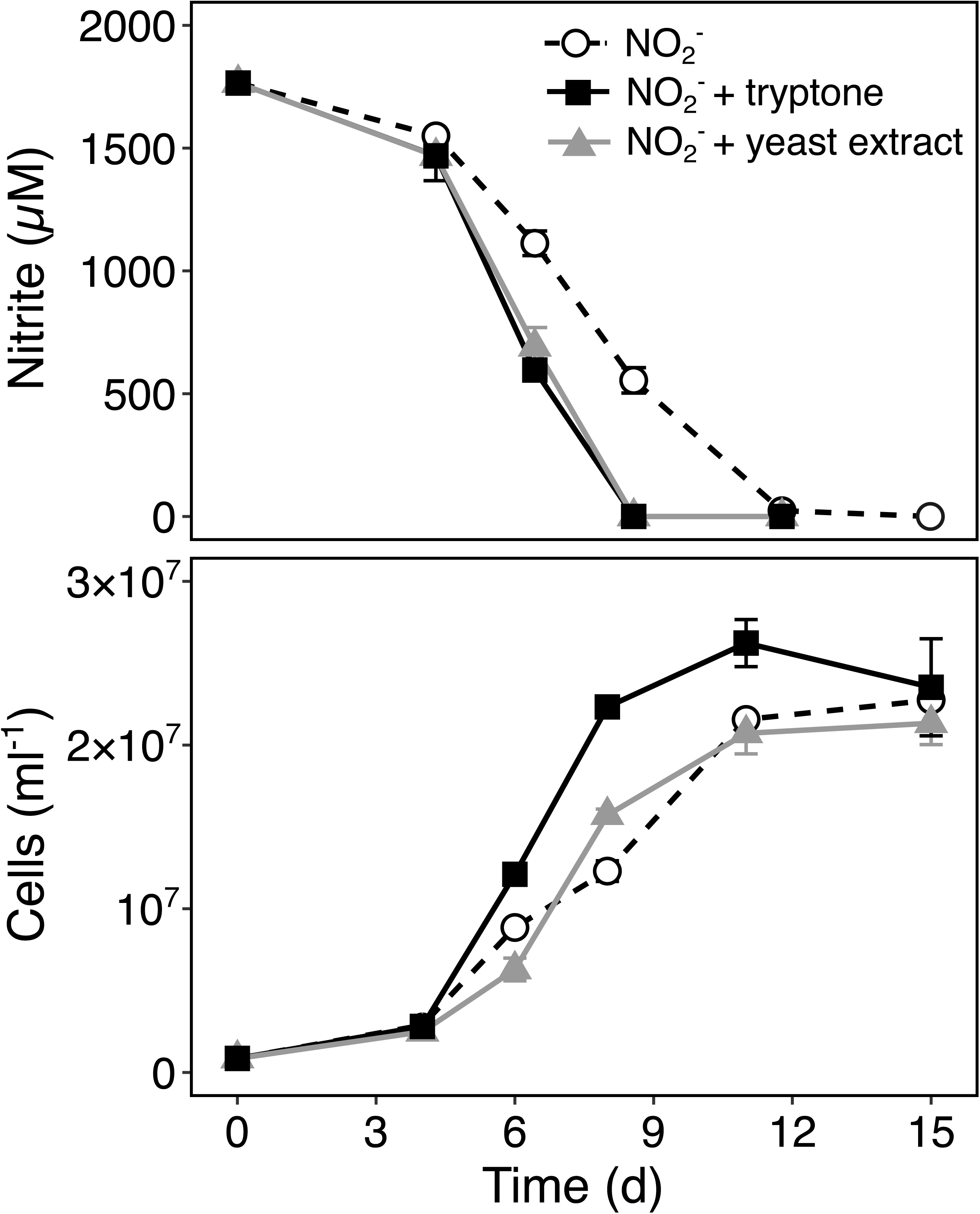
The effect of undefined organic carbon substrates on nitrite oxidation activity and growth of *N. marina* Nb-295. Error bars represent the range of measurements from duplicate cultures.

**Fig 3.**
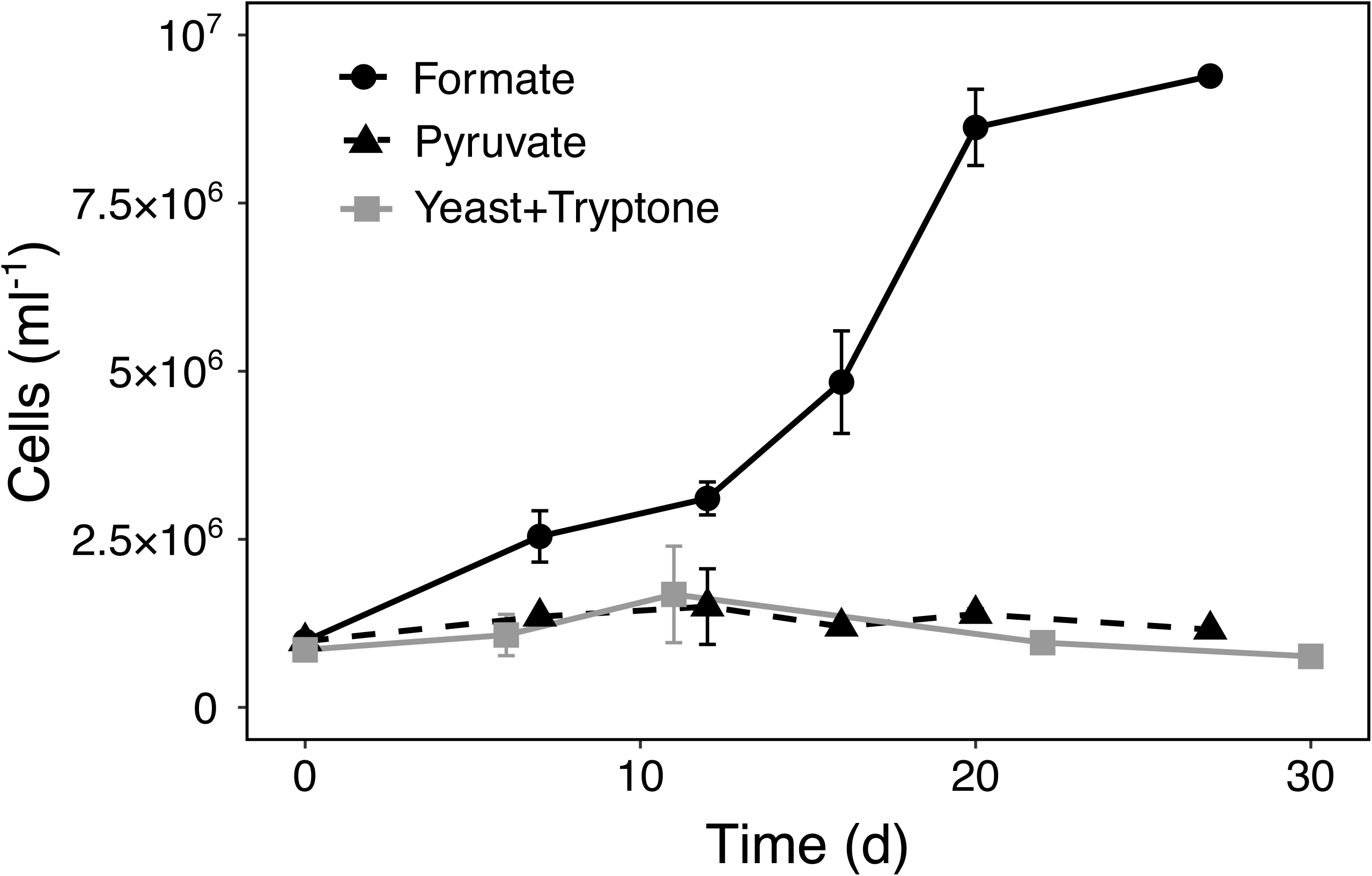
Growth of *N. marina* Nb-295 on formate (1mM), pyruvate (1mM) and yeast+tryptone (150 mg L^-1^ each) in the absence of nitrite. Error bars represent the range of measurements from duplicate cultures.

In addition to undefined organic substrates, defined organic compounds such as glycerol and pyruvate have been reported to enhance the growth of *N. marina* and *Ca*. N. defluvii, respectively [5, 12]. Furthermore, formate has been shown to serve as electron donor and carbon source for some lineage I and II *Nitrospira* [11, 25]. *N. marina* encodes a putative formate dehydrogenase (FdhA) (Table S5), which is divergent from those found in *N. moscoviensis* and *Ca*. N. defluvii (∼24 and 27% amino acid identity, respectively), but shares a relatively high sequence similarity (∼48% amino acid identity) to the functionally characterized formate dehydrogenase Fdh4 from *Methylobacterium extorquens* [63]. *N. marina* was able to grow chemoorganotrophically on 1 mM formate in the absence of NO_2_^-^ (Fig. 3), suggesting that it is not an obligate chemolithotrophic organism. However, the use of formate instead of NO_2_^-^ as electron donor resulted in slower growth rates as previously shown for *N. moscoviensis* [25]. In contrast to earlier reports [5], additions of glycerol did not stimulate NO_2_^-^ oxidation activity or growth (Table S6). Furthermore, pyruvate could neither be used as alternative energy source (Fig. 3), nor did it stimulate metabolic activity in the presence of NO_2_^-^ (Table S6). We thus hypothesize that the stimulating effect of pyruvate on *Ca*. N. defluvii could potentially be attributed to H_2_O_2_ detoxification as shown for AOA that lack catalases [45, 64].

#### Oxidative stress defense and osmoregulation

The formation of reactive oxygen species (ROS) is prevalent in oxic environments and oxidative stress defense is an important component of the stress response in marine organisms [65]. *N. marina* encodes multiple enzymes to reduce oxidative stress, including a cytoplasmic Mn/Fe-binding superoxide dismutase, a periplasmic Cu/Zn-binding SOD, two heme-containing catalases (of which KatE contains a signal peptide suggesting secretion into the periplasm or extracellular space) and various peroxiredoxins (Table S5). In contrast, *Nitrospina gracilis* and marine ammonia-oxidizing archaea lack catalase [45, 55], suggesting that *N. marina* is less susceptible to oxidative stress compared to other marine nitrifiers. In addition to its plethora of oxidative stress defense-related proteins, *N. marina* also encodes two putative photolyases, enzymes known to be involved in the repair of UV induced DNA damage [66], suggesting that it is well adapted to conditions characteristic for euphotic environments.

Marine microorganisms need to counteract the external osmotic stress from high salt concentrations by accumulating a variety of small molecules in the cytoplasm. These organic solutes (=osmolytes) can either be synthesized de-novo or transported into the cell from the surrounding environment [67]. In addition to select amino acids that can serve as compatible solutes (e.g. proline and glutamate) [68], biosynthesis pathways for the osmolytes glycine betaine and trehalose were identified in the *N. marina* genome (Table S5). *N. marina* is able to synthesize trehalose via two different pathways (either directly from maltose using trehalose synthase or from glycogen via maltodextrin and maltooligosyl-trehalose) and encodes two putative glycine betaine/osmolyte transport systems (ABC and BCCT family) to import exogenous glycine betaine (Table S5). Furthermore, despite their diverse roles and intracellular functions, polyamines can also contribute to osmotic stress protection [69, 70]. *N. marina* can synthesize the polyamines putrescine and spermidine, and encodes a putative polyamine ABC transport module (Table S5). Protection against hypo-osmotic shock in *N. marina* might also be conferred by one large- and seven small-conductance mechanosensitive ion channels, Na^+^:H^+^ antiporters and voltage-dependent K^+^ and Na^+^ channels (Table S5). The production and concomitant release of osmolytes (i.e. via diffusion, excretion, predation or cell lysis) could potentially fuel heterotrophic metabolism in the ocean [71] representing a link between chemolithoautotrophic production and heterotrophic consumption of DOM as recently suggested for ammonia-oxidizing archaea [72].

#### Vitamin biosynthesis and metal acquisition

B vitamins are important biochemical co-factors required for cellular metabolism and their concentrations are depleted to near zero in large areas of the global ocean [73]. *N. marina* encodes the complete biosynthetic pathways for the B vitamins thiamin (B_1_), riboflavin (B_2_), pantothenate (B_5_), pyridoxine (B_6_), biotin (B_7_) and tetrahydrofolate (B_9_). An incomplete cobalamin (vitamin B_12_) biosynthesis pathway was also identified, lacking genes for multiple precorrin conversion reactions that ultimately lead to the biosynthesis of the molecule’s corrin ring [74]. However, *N. marina* encodes for a putative bifunctional precorrin-2 dehydrogenase/sirohydrochlorine ferrochelatase converting precorrin-2 to siroheme, a cofactor at the active sites of many enzymes, including assimilatory nitrite reductases [75]. Since *N. marina* only encodes the cobalamin-dependent versions of methionine synthase, ribonucleotide reductase and methylmalonyl-CoA mutase, it must rely on the supply of cobalamin or its precursors by other members of the microbial community. This auxotrophy may explain why *N. marina* has previously been difficult to cultivate in a mineral medium [5]. Interestingly, the *N. marina* genome contains genes for multiple reactions that convert the precursor cobyrinate/hydrogenobyrinate to cobalamin, and encodes all genes required for cobalamin salvage from cobinamide (Table S5). Hence, in contrast to many other bacteria that lack the complete cobalamin biosynthesis pathway [76], *N. marina* appears to obtain its B_12_ solely from salvage of multiple intermediates from the environment. Cobalamin uptake is likely mediated by a TonB-dependent receptor and an ABC-type cobalamin/Fe^3+^ siderophore uptake system co-localized with cobalamin biosynthesis and salvage genes (Table S5). This absolute B_12_ requirement for three enzymes coupled to an incomplete biosynthetic pathway implies that B_12_ nutrition is an important aspect for the physiology of nitrite oxidizers. When *N. marina* was grown in natural seawater, cellular concentrations of ribonucleotide reductase were approximately 39-fold higher compared to the cyanobacterium *Prochlorococcus* [79] (Table 1) despite only having a ∼two-fold greater cell volume, consistent with the importance of B_12_ nutrition to *N. marina*. Moreover, this salvage acquisition mode is interesting in the context of dissolved cobalt speciation that is complexed by strong organic ligands hypothesized to also be B_12_ precursors or degradation products based on their thermodynamic properties [77]. In the Northwest Atlantic near where strain Nb-295 was isolated organic cobalt complexes are abundant, comprising about half the dissolved cobalt inventory [78].

In addition to B vitamins, low concentrations of iron limit ocean productivity in many parts of the global ocean [80, 81] and could limit NOB due to the high iron requirements of their respiratory chain [41, 82]. *N. marina* encodes multiple metal transport systems, including a molybdenum ABC transporter module, two putative zinc transporters (ZIP family), a putative Fe^2+^/Mn^2+^ transporter (VIT1/CCC1 family), two putative copper-transporting P-type ATPases, and a putative high-affinity nickel transport protein (Table S5), however, it lacks an ABC-type uptake system for Fe^3+^ present in other characterized *Nitrospira* species [24]. Furthermore, no known proteins associated with siderophore production were identified in the *N. marina* genome. However, the presence of multiple TonB-like receptors suggests the potential for the uptake of iron-bound-siderophores produced by other microbes. Additionally, the *N. marina* genome contains three putative bacterioferritin-related proteins for iron storage. While only one complete ABC-type cobalamin/Fe^3+^ siderophore uptake system that likely transports cobalamin (discussed above) was identified in the *N. marina* genome, an additional periplasmic substrate binding protein of a putative ABC-type cobalamin/Fe^3+^-siderophore transport system is located in proximity to iron uptake-related genes including a ferric uptake regulator (Fur) and a ferritin-like protein (Table S5). Additionally, some major facilitator superfamily-type permeases have been shown to promote bacterial iron-siderophore import from the periplasm into the cytoplasm [83]. Alternatively, iron might be released from the siderophore by a reduction process in the periplasm and subsequently imported into the cytoplasm by one of the divalent cation transporters or the putative VIT1/CCC1 family Fe^2+^/Mn^2+^ transporter, which would enable re-utilization of intact siderophores [83].

### Metabolic response to low oxygen concentrations

Given the presence of multiple signatures of microaerophilic adaptation and metabolic diversity of nitrite oxidizers [24, 25, 55, 59] and their occurrence in low oxygen environments [23, 84, 85], we sought to further investigate potential adaptations of *Nitrospira marina* Nb-295 to low O_2_ conditions. *N. marina* was grown at O_2_ concentrations characteristic for most parts of the oxic ocean (∼200 µM) and at O_2_-limiting conditions (∼5.6 µM O_2_) found in environments with high NO_2_^-^ concentrations such as oxygen minimum zones or sediments [86, 87].

When grown under atmospheric O_2_ concentration, *N. marina* oxidized 1.5 mM NO_2_^-^ within 12 days whereas NO_2_^-^ oxidation activity was decreased under O_2_-limiting conditions and depletion of 1.5 mM substrate took 27 days (Fig. 4). This result is in agreement with the initial description by Watson *et al*. [5], who reported partial inhibition at low O_2_ partial pressure. Cell abundances at the final timepoint (after 1.5 mM NO_2_^-^ was oxidized) were comparable for both treatments (Fig. S2), indicating that the reduced NO_2_^-^ oxidation activity during O_2_-limiting conditions ultimately resulted in similar cell yields.

**Fig 4.**
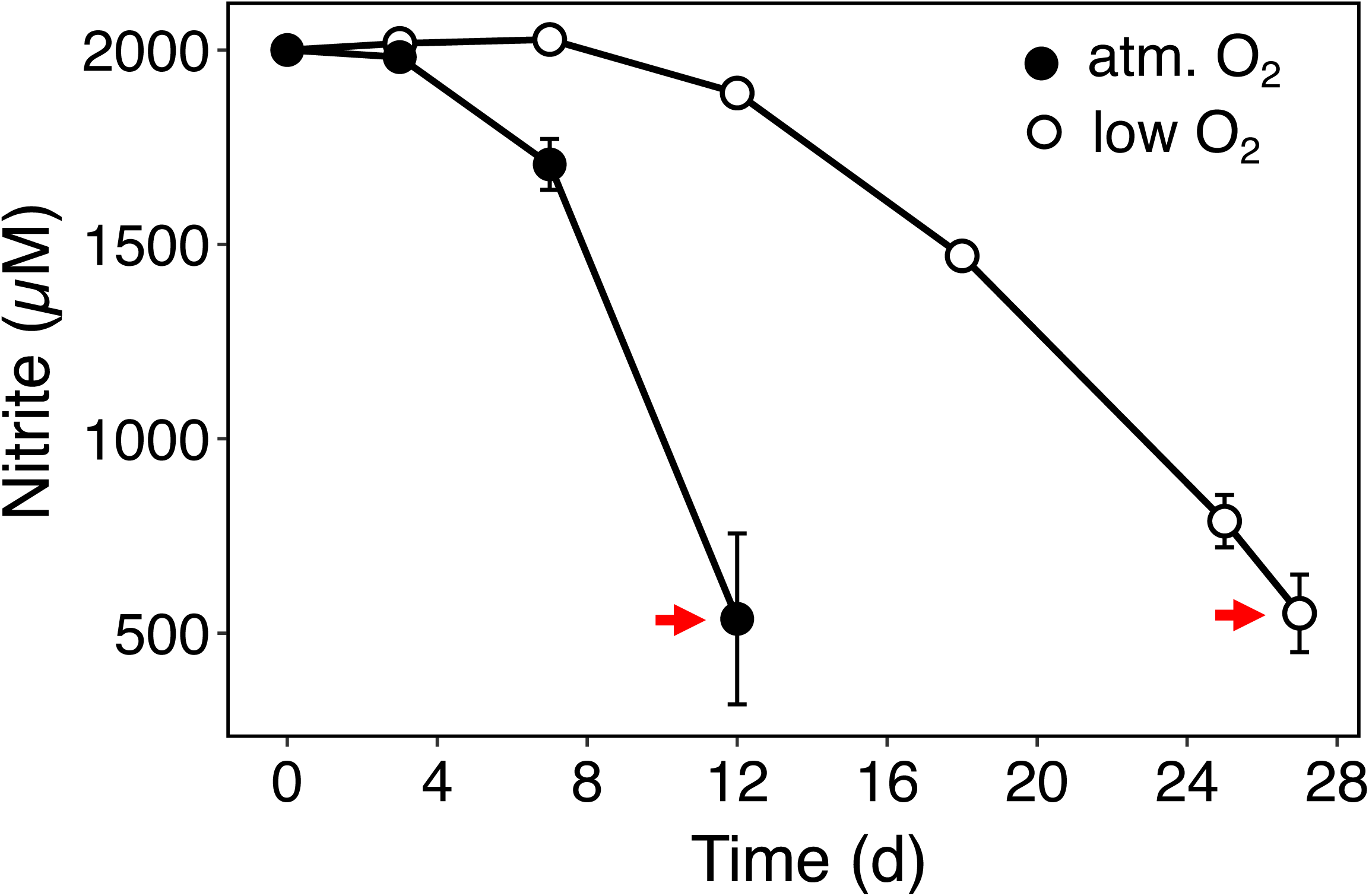
Nitrite consumption of *N. marina* Nb-295 grown under atmospheric (filled circles) and low O_2_ conditions (open circles). Cells for proteome analysis were harvested after 12 and 27 days (indicated by red arrows), respectively. Error bars represent standard deviations from measurements of triplicate cultures.

#### General proteomic response and upregulation of gene clusters

Cultures grown under atmospheric O_2_ tension and under low O_2_ concentration were harvested for proteomic analysis during exponential growth (see Material and Methods). A total of 2,086 and 2,100 proteins were identified by liquid chromatography-tandem mass spectrometry (LC-MS/MS) in the atmospheric and low O_2_ treatments, respectively, accounting for 48.8 and 49.1 % (49.9 % combined from a total of 175,653 peptides at a false discovery rate of 4.6%) of the predicted protein coding sequences (CDS) in the *N. marina* genome. As previously reported for *N. marina* and *Nitrococcus mobilis*, proteins exhibiting the highest abundances were associated with NO_2_^-^ oxidation [41] (Table S5). NXR made up on average 4% of all peptide spectral counts and cellular NXR concentrations were ∼13,500 copies cell^-1^ (Table 1), approximately covering one-fourth of the membrane surface (see Material and Methods). All three NxrA copies were detected in the proteome as determined by detection of unique peptides in each, and NxrA_3 appeared to be more abundant compared to NxrA_1 and NxrA_2 (Table 1). Under low O_2_ conditions, NxrA_1 increased in abundance (Table 1, Fig. 5) indicating different metabolic or regulatory roles of the highly similar subunits.

**Fig 5.**
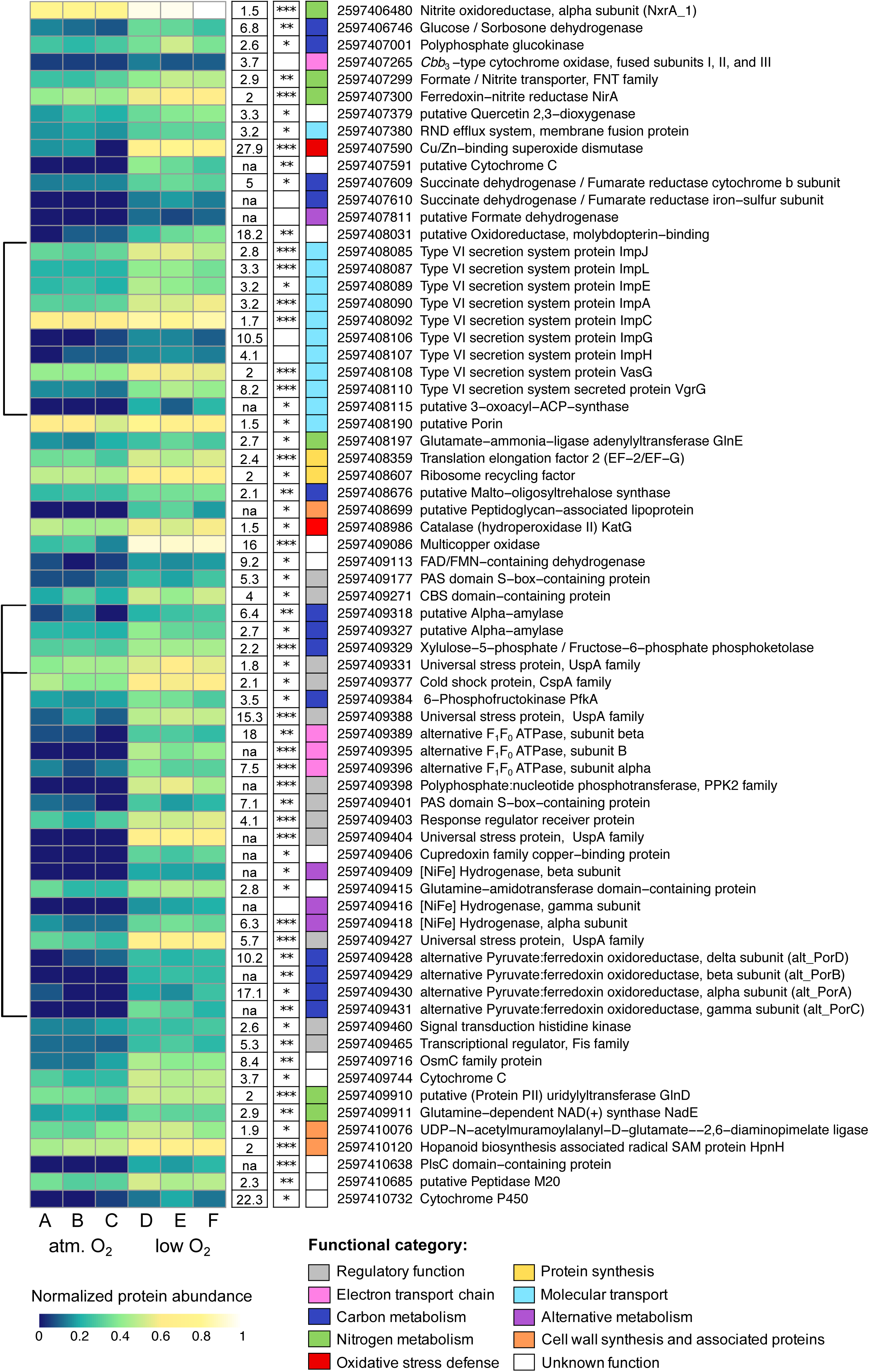
Heat map of *N. marina* Nb-295 proteins that were more abundant under low O_2_ concentrations compared to atmospheric O_2_ concentration. Relative protein abundance values were square-root transformed and hypothetical proteins were excluded to improve readability. The complete set of untransformed values can be found in Table S3. Fold-changes and significance values (adj. P value ≤ 0.001, ***; ≤ 0.01, **; ≤ 0.05, *) are shown in white boxes next to the corresponding protein. Select low abundant proteins of interest with high Fold-changes were included despite being not statistically significant (see Material and Methods). Fold-changes of proteins that were not detected under atmospheric O_2_ conditions are omitted to avoid dividing by zero (not available, na). Functional categories of depicted proteins are indicated by different colors. Gene clusters are indicated by black brackets.

Proteins involved in CO_2_ fixation, DNA replication, electron transport and central carbon metabolism were also highly abundant under both conditions, indicating that *N. marina* retained its central metabolic capacities during O_2_ deficiency (Table S5). Although the relative abundances of the majority of proteins remained constant during both treatments, 93 proteins significantly increased in abundance (adjusted P value ≤0.05) during growth at low O_2_ concentrations (Fig. 5, Table S3). These results are supported by the targeted quantitative proteomics analysis (Fig. S3, Table S2), suggesting that differences in spectral counts are a good proxy for changes in absolute protein abundances in our dataset.

Multiple universal stress proteins (UspA superfamily) were among the proteins that showed the highest increase in abundance under low O_2_ conditions compared to the control treatment (Fig. 5, Table S3). UspA proteins have versatile regulatory and protective functions to enable survival under diverse external stresses [88] and are induced during growth inhibition [89]. They are also induced in response to oxygen starvation in *Mycobacterium smegmatis* [90] and contribute to the survival of *Pseudomonas aeruginosa* under anaerobic conditions [91]. In *N. marina*, all four upregulated UspA proteins are located upstream or downstream of operons containing genes that also increased in abundance under low O_2_ conditions (Fig. 5), suggesting a regulatory role of UspA-related proteins upon O_2_ limitation. These upregulated gene clusters include proteins involved in electron transport, carbon metabolism and alternative energy metabolism (Fig. 5).

In addition to putative UspA-regulated gene clusters, a gene cluster containing type VI secretion system (T6SS)-related proteins exhibited higher abundances under low O_2_ conditions (Fig. 5). The T6SS is typically involved in the secretion of effectors required for pathogenesis, bacterial competition, biofilm formation, and cell communication (e.g., quorum sensing) [92, 93]. While quorum sensing has recently been shown for diverse NOB including *Nitrospira moscoviensis* [94], no LuxI autoinducer synthases and/or LuxR signal receptor homologues were identified in the *N. marina* genome. Hence, the role of T6SS in *N. marina* remains to be determined.

#### Induction of a putative O_2_-sensitive 2-oxoacid:ferredoxin oxidoreductase complex

The key enzymes of the rTCA cycle, pyruvate:ferredoxin oxidoreductase (POR) and 2-oxoglutarate:ferredoxin oxidoreductase (OGOR), are typically highly O_2_ sensitive because they contain easily oxidized iron-sulfur clusters [95]. In *Hydrogenobacter thermophilus*, five-subunit O_2_-tolerant forms of POR and OGOR mainly support aerobic growth, while a O_2_-sensitive two-subunit form is used under anaerobic conditions [96]. *N. marina* encodes three 2-oxoacid:ferredoxin oxidoreductase gene clusters that could exhibit POR or OGOR activity (Table S5). Two of these gene clusters consist of five CDS which exhibit a high sequence similarity to the O_2_-tolerant five-subunit POR/OGOR of *H. thermophilus* [97, 98], as previously described for *Ca*. N. defluvii [24]. Both complexes were highly abundant in *N. marina* proteomes from atmospheric and low O_2_ treatments (Table S5, Table 1) confirming their important role in central carbon metabolism. The third cluster contains alpha, beta and gamma subunits of a putative POR with homology to the functionally characterized four-subunit PORs of anaerobic thermophiles *Pyrococcus* and *Thermotoga* [99] and is absent in all other *Nitrospira* with the exception of *Ca*. N. alkalitolerans [9]. A protein with a 4Fe-4S binding domain was identified in the same operon, potentially representing the missing delta subunit of the POR complex (Table S5). This putative four-subunit POR was among the proteins that showed the highest increase in abundance under low O_2_ conditions in *N. marina* (Fig. 5). The O_2_-tolerant POR/OGOR isoforms were reported to have a >5 times lower specific activity [96] and might therefore constitute a substantial part of the cellular soluble protein content in *H. thermophilus* [100]. Hence, it is tempting to speculate that *N. marina* increases the expression of a more efficient (i.e. higher specific activity), O_2_-sensitive four-subunit POR under O_2_ limitation. While oxidative stress typically decreases under low O_2_ conditions, it might still be high enough to damage O_2_-sensitive enzymes. *In N. marina*, the abundance a periplasmic Cu/Zn-binding superoxide dismutase (SOD) and a cytoplasmic catalase (KatG) increased under O_2_-limited conditions (Fig. 5), while the abundances of other oxidative stress defense-related proteins remained constant (Table S5). SOD has been shown to be efficient in protecting POR activity from oxidative damage in *Entamoeba histolytica* [101]. In *N. marina*, SOD was among the proteins with the highest increase in abundance under low O_2_ conditions (foldchange: 27.9), suggesting a role in POR protection.

#### Expression of a high O_2_-affinity cbb_3_-type terminal oxidase

The majority of proteins related to electron transport showed similar abundance levels during growth at atmospheric and limiting O_2_ concentrations (Table S5). Despite its overall low abundance, a putative high-affinity cytochrome *cbb*_*3*_-type terminal oxidase was 3.6-times more abundant under low O_2_ concentrations compared to atmospheric O_2_ tension (Fig. 5). The *cbb*_*3*_-type terminal oxidase of *Bradyrhizobium japonicum* was reported to have a K_m_ value of 7 nmol L^-1^ O_2_ [102] and NO_2_^-^ oxidation rates have been detected at O_2_ concentrations in the low nanomolar range (5-33 nmol L^-1^ O_2_) [23]. This suggests that the *cbb*_*3*_-type terminal oxidase might enable *N. marina* to retain aerobic respiration at low O_2_ concentrations, albeit at lower NO_2_^-^ oxidation rates (Fig. 4). Low-affinity terminal oxidases are typically more efficient in energy conservation [103], indicating that *N. marina* benefits from the presence of a terminal oxidase with lower O_2_ affinity in well-oxygenated environments. While *N. gracilis* encodes a highly similar high affinity *cbb*_*3*_-type terminal oxidase as *N. marina* [55], this enzyme is lacking in most *Nitrospinae*, including those identified in oxygen minimum zones (OMZ) and sediments [85, 104, 105]. Interestingly, these genomes also lack the putative terminal oxidase proposed for *Nitrospira* [24]. Still, in an OMZ where *Nitrospinae* bacteria were the only detected NOB, NO_2_^-^ oxidation rates already approached saturation at ∼1 μmol L^-1^ O_2_ [23], indicating that *Nitrospinae* bacteria might be better adapted to low O_2_ concentrations compared to *N. marina*, but the enzyme conferring high O_2_ affinity in *Nitrospinae* remains to be identified.

#### Upregulation of genes involved in alternative energy metabolism

The abundance of proteins putatively involved in alternative energy metabolisms increased under low O_2_ concentrations. These included type 3b [NiFe] hydrogenase and formate dehydrogenase (Fig. 5). Alternative electron donors such as H_2_ and formate are common reaction products at oxic/anoxic interfaces [106]. *N. moscoviensis* can couple H_2_ and formate oxidation to NO_3_^-^ reduction to remain active under anoxia [25, 27] (whereas no net growth was observed for the latter [25]), however, it is unlikely that H_2_ or formate were present at the culture conditions in this study. While abundances of hydrogenase and formate dehydrogenase increased under low O_2_ conditions, they were overall still comparably low (Fig. 5), suggesting that their expression might be regulated when conditions become more unfavorable for aerobic NO_2_^−^ oxidation. Curiously, multiple subunits of a Na^+^-translocating ATPase (alternative complex V) exhibited higher abundances at low O_2_ concentration (Fig. 5). Na^+^-translocating ATPases are suggested to be ancient enzymes that were later replaced by energetically more favorable H^+^-translocating ATPases [107]. While only few obligate anaerobes with very tight energy budgets (which cannot cover the losses caused by proton leaks) primarily use Na^+^ energetics, many organisms retained Na^+^ pumps and utilize them under energetically less favorable conditions such as anaerobiosis [107]. Hence, expression of a Na^+^-translocating ATPase suggests an adaptation of *N. marina* to overcome periods of starvation when energetically favorable electron donors or acceptors are short in supply.

## Conclusions

Although the vast majority of the ocean is well oxygenated, oxygen-depleted zones exist within the oceanic water column and in marine sediments [86, 87] with consequences for microbial adaptation and evolution. Our results show that, in contrast to some marine NOB populations, NO_2_^-^ oxidation activity of *Nitrospira marina* was reduced when grown at ∼5.6 µM L^-1^ O_2_, suggesting different O_2_ adaptations among different NOB. In accordance with its faster growth rates at atmospheric O_2_ tension, *N. marina* encodes a plethora of proteins involved in oxidative stress defense. We confirm that *N. marina* benefits from the addition of undefined organic substrates, which were shown to be inhibitory for *Nitrospina gracilis* [108], potentially further contributing to ecological niche partitioning within marine NOB. Our results indicate that *N. marina* is highly metabolically versatile, which might enable it to survive under unfavorable conditions with fluctuating levels of electron donors and acceptors. Hence, while *Nitrospinae* bacteria are the dominant nitrite oxidizers in oligotrophic oceanic regions and OMZs, *Nitrospira* might be better adapted to well-oxygenated high productivity regions including coastal systems, deep-sea trenches and hydrothermal vents. Finally, our results also indicate several previously unreported ways that NOB may interact with other members of the marine microbial community—through the supply of organic carbon-containing osmolytes and their requirement for exogenous cobalamin.

## Acknowledgements

We thank John B. Waterbury and Frederica Valois for providing the culture of *Nitrospira marina* Nb-295^T^. The *N. marina* genome was sequenced as part of US Department of Energy Joint Genome Institute Community Sequencing Project 1337 to CLD, AES, and MAS in collaboration with the user community. We thank Claus Pelikan for bioinformatic assistance. This research was supported by a Simons Foundation Early Career Investigator in Marine Microbiology and Evolution Award (345889) and US National Science Foundation (NSF) award OCE-1924512 to AES. Proteomics analysis was supported by NSF awards OCE-1924554 and OCE-1850719, and NIH award 1R01GM135709-01A1 to MAS. BB was supported by the Austrian Science Fund (FWF) Project Number: J4426-B (“The influence of nitrifiers on the oceanic carbon cycle”), SL by the Netherlands Organization for Scientific Research (NWO) grant 016.Vidi.189.050, and CLD by NSF award OCE-125999.

## Data availability

The genome of *Nitrospira marina* Nb-295^T^ is available in the JGI IMG/M repository under genome ID number 2596583682. Targeted peptide concentrations are available in Supplementary Table 2, global proteomic spectral counts and differential expression analysis are available in Supplementary Table 3. Raw mass spectra are available in PRIDE as project number XXXX.

