## Supplementary Material for "Metabolic versatility of the nitrite-oxidizing bacterium *Nitrospira marina* and its proteomic response to oxygen-limited conditions"

### Supplementary Information (SI)

#### Supplementary Methods

*Flow cytometry cell counts*

Samples of 500 µl were collected in 2 mL sterile polypropylene tubes, fixed with glutaraldehyde (0.5% final concentration) for 10 min and subsequently frozen and stored at -80°C. Prior to analysis, samples were diluted (1:10, 1:50 or 1:100, depending on the cell concentration) in 0.2 µm filtered Tris-EDTA buffer (1M Tris-HCl, 0.1M EDTA, pH 8) and stained with SYBR Green I (Invitrogen) at a final concentration of 1:5,000 for 30 min. Enumeration of cells was performed on an Easy-Cyte flow cytometer (Guava Technologies).

Protein extraction and purification

Samples were resuspended with 1800 $\mu$l of 1% SDS extraction buffer (1% SDS, 0.1M Tris/HCl pH 7.5, 10mM EDTA). Each sample was incubated at room temperature for 15 minutes, heated at 95°C for 10 minutes, and shaken at room temperature (RT) at 350 rpm for 1 h. The protein extracts were decanted and centrifuged at 14100 x g for 20 min at RT. The supernatants were removed and concentrated by membrane centrifugation to approximately 300 $\mu$l in 5 K MWCO Vivaspin units (Sartorius Stedim, Goettingen, Germany). Each sample was precipitated with cold 50% methanol/50% acetone/0.5 mM HCl for 3 days at –20°C, centrifuged at 14100 x g for 30 min at 4°C, decanted and dried by vacuum concentration (Thermo Savant Speedvac) for 10 min or until dry. Pellets were resuspended in 1% SDS extraction buffer and left at RT for 1 h to completely dissolve. Total protein was quantified (Bio-Rad DC protein assay, Hercules, CA) with BSA as a standard.

Extracted proteins were purified from SDS detergent, reduced, alkylated and trypsin digested while embedded within a polyacrylamide tube gel, modified from a previously published method [1]. A gel premix was made by combining 1 M Tris HCl (pH 7.5) and 40% Bis-acrylimide L 29:1 (Acros Organics) at a ratio of 1:3. The premix (103 $\mu$l) was combined with an extracted protein sample (35 $\mu$g-50 $\mu$g), TE Buffer, 7 $\mu$l 1% APS and 3 $\mu$l of TEMED (Acros Organics) to a final volume of 200 $\mu$l. After 1 h of polymerization at RT, 200 $\mu$l of gel fix solution (50% ETOH, 10% acetic acid in LC/MS grade water) was added to the top of the gel and incubated at room temperature for 20 minutes. Liquid was then removed and the tube gel was transferred into a new

1.5 ml microtube containing 1.2 ml of gel fix solution then incubated at RT, 350 rpm in a Thermomixer R (Eppendorf) for 1 h. The gel fix solution was then removed and replaced with 1.2 ml destaining solution (50% MeOH, 10% acetic acid in $H_2O$) and incubated at 350 rpm for 2 h. Liquid was then removed, gel cut up into 1 mm cubes and then added back to tubes containing 1 ml of 50:50 acetonitrile:25 mM ammonium bicarbonate (ambic) incubated for 1 h, 350 rpm at RT. Liquid was removed and replaced with fresh 50:50 acetonitrile:ambic and incubated at 16°C at 350 rpm overnight. The above step was repeated for 1 hour the following morning. Gel pieces were then dehydrated twice in 800 $\mu$l of acetonitrile for 10 min at room temperature and dried for 10 min in a ThermoSavant DNA110 speedvac after removing solvent. 600 $\mu$l of 10 mM DTT in 25 mM ambic was added to reduce proteins incubating at 56°C, 350 rpm for 1 h. Unabsorbed DTT solution was then removed with volume measured. Gel pieces were washed with 25 mM ambic and 600 $\mu$l of 55 mM iodacetamide was added to alkylate proteins at RT, 350 rpm for 1 h. Gel cubes were then washed with 1 mL ambic for 20 min, 350 rpm at RT. Acetonitrile dehydrations and speedvac drying were repeated as above. Trypsin (Promega #V5280) was added in appropriate volume of 25 mM ambic to rehydrate and submerse gel pieces at a concentration of 1:20 $\mu$g trypsin:protein. Proteins were digested overnight at 350 rpm and 37°C. Unabsorbed solution was removed and transferred to a new tube. 50 $\mu$l of peptide extraction buffer (50% acetonitrile, 5% formic acid in water) was added to gels, incubated for 20 min at RT then centrifuged at 14,100 x g for 2 min. Supernatant was collected and combined with unabsorbed solution. The above peptide extraction step was repeated combining all supernatants. Combined protein extracts were centrifuged at 14,100 x g for 20 minutes, supernatants transferred into a new tube and dehydrated down to approximately 10-20 $\mu$l in the speedvac. Concentrated peptides were then diluted in 2% acetonitrile 0.1% formic acid in water for storage until analysis. All water used in the tube gel digestion protocol was LC/MS grade, and all plastic microtubes were ethanol rinsed and dried prior to use.

*Phylogenetic analyses*

Phylogenomic analysis was performed using 120 concatenated phylogenetic marker genes of representatives of the phylum *Nitrospirae/Nitrospirota* as implemented in the Genome Taxonomy Database Toolkit (GTDB-tk) version 1.1.1. [2]. The multiple sequence alignment contained 95 genomes and metagenome-assemble genomes (MAGs) from the Genome Taxonomy Database (GTDB) (Release 04-RS89, 19th June 2019) [3], the genome of *Nitrospira marina* Nb-295$^T$ (this study), the MAG of *Ca.* Nitrospira alkalitolerans [4] and two open ocean single-amplified genomes

(SAGs) AC-738-G23 and AC-732-L14 [5]. All genomes were estimated to be ≥50% complete with ≤5% contamination based on CheckM [6]. Conserved marker genes were identified using the command 'gtdbtk identify' (default parameters) and aligned to the reference genome alignment using the command 'gtdbtk align' using a taxonomy filter for *Nitrospirae/Nitrospirota* (--taxa_filter p__Nitrospirota,p__Nitrospirota_A) as implemented in GTDB-Tk. The resulting alignment was used to calculate a maximum likelihood phylogenetic tree with IQ-Tree version 1.6.9 [7] based on the best-fit model (LG+F+R8), with ultrafast bootstrap (UFBoot) inferred from 10,000 replicates. The phylogenetic tree was visualized with the online tool iTOL [8].

Full-length 16S rRNA gene sequences were obtained from the GenBank database at the National Center for Biotechnology Information (NCBI) [9], except for *Nitrospira marina* Nb-295$^T$ and the SAGs AC-728-G23 and AC-728-O15, which were obtained from the Integrated Microbial Genomes and Microbiomes (IMG) platform [10], and the sequence of *Ca.* Nitrosospira alkalitolerans KS, which was obtained from the MicroScope platform [11]. Gene sequences were aligned with MAFFT (L-INS-I method) [12] and unreliable positions were filtered from the resulting alignments with BMGE [13] resulting in 1361 nucleotide positions in the final alignment (except for the partial sequence of *N. bockiana* which only contained 1095 nucleotide positions). The phylogenetic tree was calculated by maximum likelihood with IQ-Tree version 1.6.9 [7] based on the best-fit model (TIM3+F+I+G4), with ultrafast bootstrap (UFBoot) inferred from 10,000 replicates. The phylogenetic tree was visualized with the online tool iTOL [8]. *Leptospirillum ferrooxidans* (X86776) was used as outgroup

#### Supplementary Figures and Tables

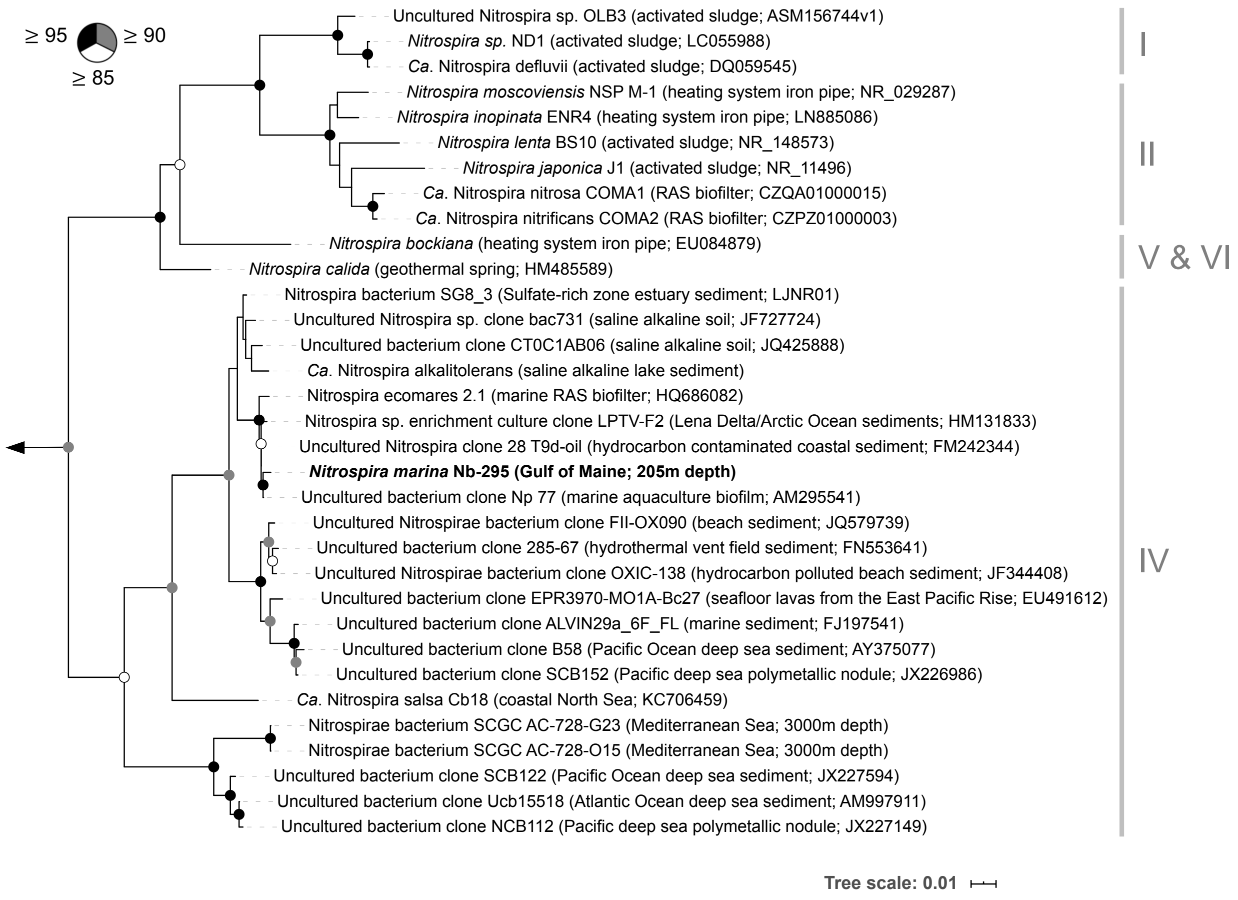


Fig S1. Maximum likelihood phylogenetic analysis of 16S rRNA gene sequences of cultured representatives and selected environmental sequences from the genus *Nitrospira*. *Leptospirillum ferrooxidans* (X86776) was used as outgroup indicated by the black arrow. Support values ≥85% are represented on the respective branches by circles color-coded as indicated in the figure. The scale bar represents 0.01 substitutions per site.

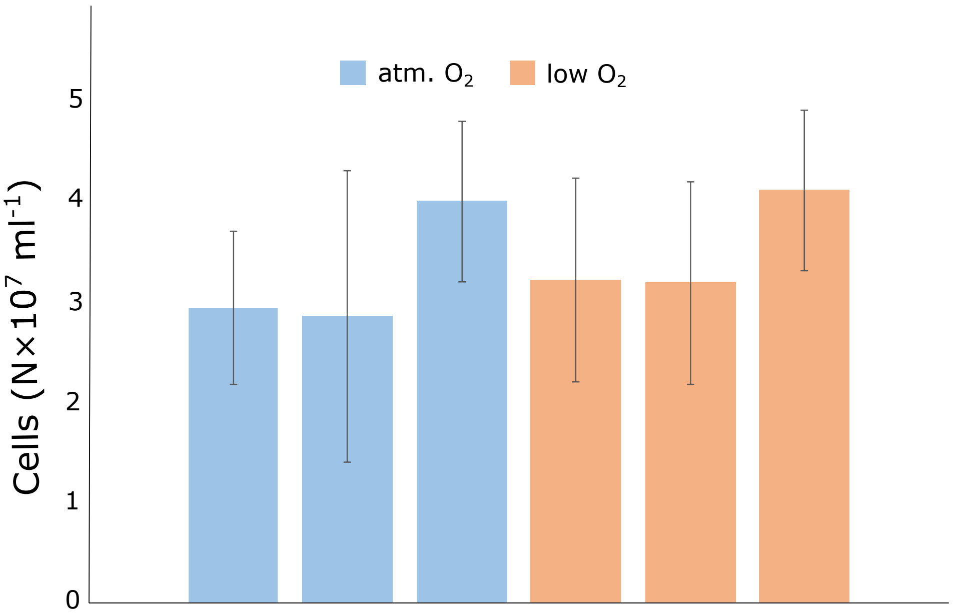


Figure S2. Cell counts of *N. marina* Nb-295 at the final time point of the oxygen experiments before cells were harvested for proteome analyses. Error bars represent standard deviations of cell counts from triplicate filters of the same sample.

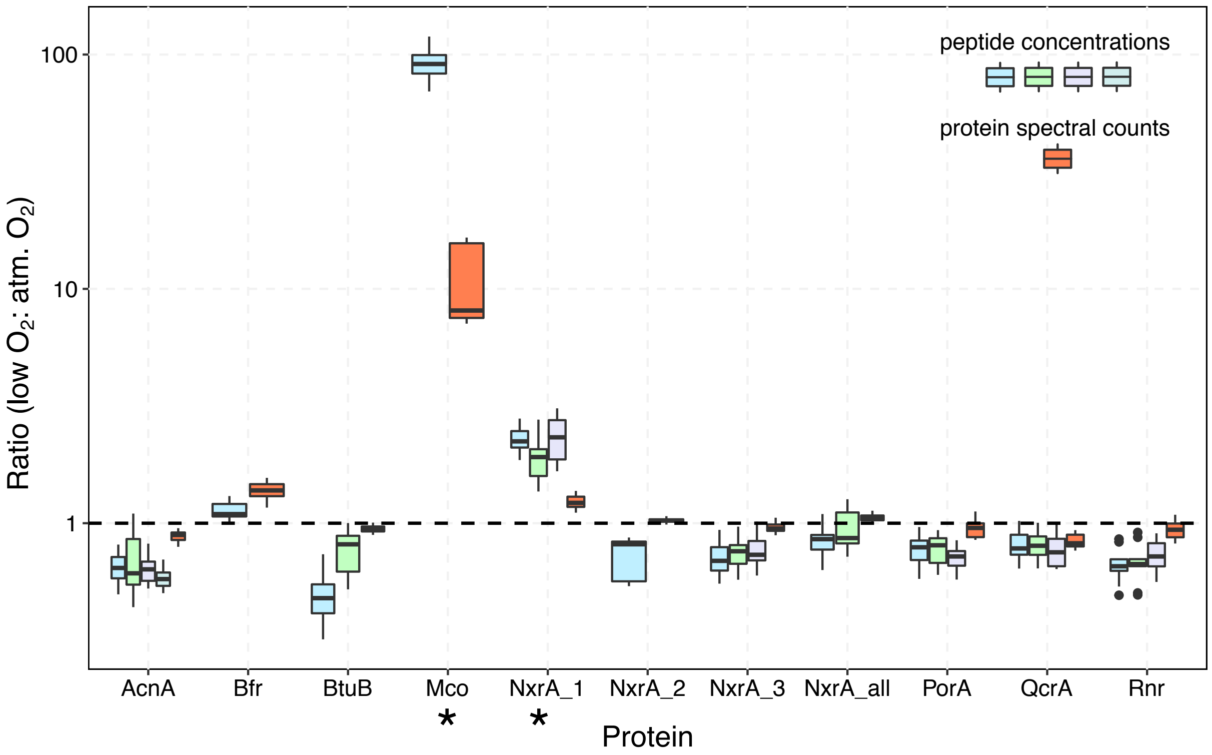


Figure S3. Comparison of proteomic spectral counts and quantified peptide concentrations. The dashed line indicates equal protein abundances in both treatments. Proteins marked with an asterisk (*) were significantly more abundant under low $O_2$ conditions (see Table S3 and Figure 5 in the main text).

Table S1. Composition of the nitrite oxidizer artificial seawater (NOASW) medium

#### Stock solutions:

| **Basic salt solution** | **1.5L** | |
|---|---|---|
| NaCl | 37.08 | g |
| $MgSO_4$ $7H_2O$ | 9.4 | g |
| $MgCl_2$ $6H_2O$ | 7.1 | g |
| $CaCl_2$ $2H_2O$ | 2.05 | g |
| KCl | 1.01 | g |
| $NaHCO_3$ | 0.27 | g |
| fill up to 2L with MilliQ | | |

| **Sodium nitrite solution (1M)** | **100** | **mL** |
|---|---|---|
| $NaNO_2$ | 6.9 | g |
| fill up to 100mL with MilliQ | | |

| **$K_2HPO_4$ solution (0.05M)** | **100** | **mL** |
|---|---|---|
| $K_2HPO_4$ | 0.87 | g |
| fill up to 100mL with MilliQ | | |

| **FeNaEDTA Solution (1g/L)** | **100** | **mL** |
|---|---|---|
| FeNaEDTA | 100 | mg |
| fill up to 100mL with MilliQ | | |

| **Modified Non-chelated trace element mixture** | | | **1L** |
|---|---|---|---|
| Distilled $H_2O$ | 987 | mL | |
| HCl (conc. ~12.5M) | 8 | mL | (100mM) |
| $H_3BO_3$ | 30 | mg | (0.5mM) |
| $MnCl_2$ $4H_2O$ | 20 | mg | (0.1mM) |
| $CoCl_2$ $6H_2O$ | 100 | mg | (0.5mM) |
| $NiCl_2$ $6H_2O$ | 24 | mg | (0.1mM) |
| $CuCl_2$ $2H_2O$ | 20 | mg | (0.1mM) |
| $ZnSO_4$ $7H_2O$ | 144 | mg | (0.5mM) |
| $Na_2MoO_4$ $2H_2O$ | 24 | mg | (0.1mM) |

| **1000x Vitamin B12 solution (1mg/L)** | **1** | **L** |
|---|---|---|
| Cyanocobalamin | 1 | mg |
| fill up to 1L with MilliQ | | |

#### Medium preparation:

| | 1L | |
|---|---|---|
| Basic salt solution | 995 | mL |
| $NaNO_2$ (1M) | 2 | mL |
| Modified Trace Elements | 1 | mL |
| FeNaEDTA Solution (1 g/L) | 0.5 | mL |
| $KH_2PO_4$ (0.05 M) | 0.5 | mL |
| 1000x Vitamin B12 solution | 1 | mL |

Table S4. Genome characteristics of *N. marina* Nb-295 and other cultured nitrite-oxidizing bacteria

| *Nitrospira* Lineage | IV | IV | I | II | II | II | II | II | II | | |
|---|---|---|---|---|---|---|---|---|---|---|---|
| **Species** | ***N. marina* Nb-295** | ***Ca.* N. alkalitolerans KS** | ***Ca.* N. defluvii** | ***N. moscoviensis* NSP M-1** | ***N. japonica* NJ1** | ***N. lenta* BS10** | ***N. inopinata* ENR4** | ***Ca.* N. nitrosa COMA1** | ***Ca.* N. nitrificans COMA2** | ***Nitrococcus mobilis* Nb-231** | ***Nitrospina gracilis* 3/211** |
| Reference | this study | Daebeler et al. 2020 | Lücker et al. 2010 | Koch et al. 2014 | Ushiki et al. 2018 | Sakoula et al. 2018 | Daims et al. 2015 | van Kessel et al. 2015 | van Kessel et al. 2015 | Füssel et al. 2017 | Lücker et al. 2013 |
| Environment | ocean | saline alkaline lake | WWTP | heating system pipe | WWTP | WWTP | heating system pipe | WWTP | WWTP | ocean | ocean |
| Genome size (Mb) | 4.68 | 4.94 | 4.32 | 4.59 | 4.08 | 3.76 | 3.30 | 4.42 | 4.12 | 3.62 | 3.07 |
| Closed genome? | yes | no | yes | yes | yes | no | yes | no | no | no | no |
| Coding DNA sequences | 4272 | 5091 | 4274 | 4508 | 4150 | 3968 | 3024 | 4309 | 4502 | 4052 | 3064 |
| Average G+C content (%) | 50 | 51.4 | 59 | 62 | 59 | 57.9 | 59.2 | 54.8 | 56.6 | 60 | 56.2 |
| Number of rRNAs | 3 | 3 | 3 | 4 | 3 | 3 | 3 | 3 | 3 | 3 | 3 |
| Number of tRNAs | 47 | 47 | 46 | 47 | 45 | 46 | 47 | 46 | 43 | 45 | 45 |
| NxrAB operons | 3 | 1[a]. | 2 | 4 (5)[b] | 3 | 2 | 1 | 2 | 4 | 1[c] | 2 |
| Urease | – | – | – | + | + | + | + | + | + | – | – |
| Cyanase | + | + | + | + | + | + | – | – | – | + | + |
| Hydrogenase | 3b | 2a, 3b | – | 2a | – | – | 3b | 3b | 3b | 3b | 3b |
| Formate dehydrogenase | + | + | + | + | – | – | – | – | – | + | – |
| Catalase | 2 | 1 | 0 | 3 | 2 | 0 | 1 | 0 | 1 | 1 | 0 |
| Superoxide dismutase | 2 | 2 | 0 | 1 | 2 | 2 | 0 | 0 | 1 | 2 | 0 |
| Flagella | + | + | + | + | – | + | + | + | + | + | + |

[a] contains one complete NxrAB operon, an additional complete NxrA copy and one fragmented NxrA copy at the end of a contig
[b] lost one NxrAB operon during prolonged cultivation
[c] contains an additional NxrA copy

Table S6. The effect of culture medium amendments on nitrite oxidation (A) and growth (B) of *N. marina* Nb-295. All incubations contained 2 mM $NaNO_2^-$ and 1X Vitamin $B_{12}$ solution (see Table S1 for medium preparation). Nitrite concentrations are given in µM (A) and cell abundances in $x10^5$ $ml^{-1}$ (B).

A

| Time (d) | Control | Ammonia (100 µM) | Pyruvate (500 µM) | Glycerol (1 g $L^{-1}$) | Yeast (150 mg $L^{-1}$) | Tryptone (150 mg $L^{-1}$) |
|---|---|---|---|---|---|---|
| 0 | 1788 ± 2.8 | 1788 ± 2.8 | 1788 ± 2.8 | 1788 ± 2.8 | 1788 ± 2.8 | 1788 ± 2.8 |
| 7 | 519.5 ± 41.7 | 457 ± 1.41 | 507.5 ± 19.1 | 508 ± 114.6 | 60.5 ± 46 | 0 ± 0 |
| 11 | 0 ± 0 | 0 ± 0 | 0 ± 0 | 0 ± 0 | 0 ± 0 | 0 ± 0 |

B

| Time (d) | Control | Ammonia (100 µM) | Pyruvate (500 µM) | Glycerol (1 g $L^{-1}$) | Yeast (150 mg $L^{-1}$) | Tryptone (150 mg $L^{-1}$) |
|---|---|---|---|---|---|---|
| 0 | 9.9 ± 0 | 9.9 ± 0 | 9.9 ± 0 | 9.9 ± 0 | 9.9 ± 0 | 9.9 ± 0 |
| 7 | 28.7 ± 0.7 | 26.5 ± 0.9 | 27.7 ± 2.1 | 30.3 ± 1 | 22.8 ± 0.7 | 38.5 ± 1.1 |
| 11 | 200.4 ± 15.2 | 202.8 ± 7.7 | 206.6 ± 9.1 | 206 ± 0.6 | 226.3 ± 36.6 | 253.2 ± 17.8 |
